# Junctophilin-2 promotes cardiomyocyte survival by blocking MURF1-mediated Junctin ubiquitination and proteasome-dependentdegradation

**DOI:** 10.1101/2022.10.23.513420

**Authors:** Xiaoyun Ji, Yifan Huang, Rui Ni, Dong Zheng, Guo-Chang Fan, Douglas L Jones, Long-Sheng Song, Subrata Chakrabarti, Zhaoliang Su, Tianqing Peng

## Abstract

**Aims:** Junctophilin-2 is required for the development, maturation and integrity of the t-tubule system and the gating stability of RyR2 in cardiomyocytes. This study investigated whether and how junctophilin-2 maintained junctin, a scaffold protein stabilizing RyR2, to prevent cardiomyocyte death under stress.

**Methods:** Cardiomyocytes were exposed to conditions of stress including palmitate, doxorubicin, or hypoxia/re-oxygenation. Adenoviral vectors were employed to manipulate expression of junctophilin-2 and junctin in cardiomyocytes. Molecular/cellular/biochemical analyses were conducted.

**Results:** Different conditions of stress decreased junctophilin-2 expression through aberrant autophagy and concomitantly induced a reduction of junctin protein in cardiomyocytes. Over-expression of junctophilin-2 preserved the protein levels of junctin and attenuated cytosolic Ca^2+^ and apoptosis in cardiomyocytes under stress. Knockdown of junctophilin-2 reproduced the detrimental phenotypes of stress in cardiomyocytes. Notably, over-expression of junctin prevented cardiomyocyte death under stress whereas knockdown of junctin offset the protective effects conferred by junctophilin-2 over-expression. Mechanistically, junctophilin-2 blocked MURF1-junctin interaction thereby preventing junctin ubiquitination and proteasome-dependent degradation. Mass spectrometry analysis identified multiple ubiquitination sites on the junctin protein and the non-ubiquitinated junctin mutant (K8A/K102A/K107A/K140A) was resistant to degradation.

**Conclusions:** This study uncovers an unrecognized role of junctophilin-2 in preventing junctin ubiquitination and degradation in maintaining cytosolic Ca^2+^ homeostasis. Both junctophilin-2 and junctin represent two new survival factors of cardiomyocytes and thus, may be new therapeutic targets for cardiac protection.

## Introduction

Junctophilin-2 (JPH2) is an important structural protein in forming junctional membrane complexes, which is essential for excitation-contraction coupling of cardiomyocytes^1^. JPH2 tethers junctional sarcoplasmic reticulum with invaginated T-tubule membrane in cardiomyocytes ^2, 3^. Additionally, JPH2 is also necessary for the development of postnatal T-tubules in mammals^4, 5^. However, JPH2 protein decreases in diseased hearts^5–13^. Loss of JPH2’s function by mutations or down-regulation of its protein expression is associated with hypertrophic cardiomyopathy, arrhythmias, and the progression of heart failure^6, 14, 15^. In contrast, over-expression of JPH2 attenuated heart failure development after cardiac stress^16^. Further studies reveal that JPH2 encodes a stress-adaptive transcriptional regulator, whose N-terminus translocates to the nucleus and represses maladaptive transcriptional reprogramming, protecting against pathological cardiac remodeling in stressed hearts^12^. Notably, recent studies have found that in addition to its interaction with the L-type Ca^2+^ channel in regulating Ca^2+^-induced Ca^2+^ release ^17, 18^, JPH2 also plays a critical role in ryanodine receptor-2 (RyR2) gating stability through preventing abnormal RyR2-mediated Ca^2+^ leaks in mature cardiomyocytes^14, 19, 20^. These previous studies have highlighted critical roles of JPH2 in cardiac physiology and pathology. However, it has never been reported if JPH2 plays a role in cardiomyocyte survival/death.

Junctin (JCN) is an accessory protein of the sarco/endoplasmic reticulum Ca^2+^ release unit that interacts with RyR2 and Calsequestrin^21^. Genetic ablation of JCN in the mouse heart reveals that JCN is essential for the maintenance of normal RyR2 activity and Ca^2+^ release and thus, JCN-deficient cardiomyocytes have excessive RyR2 spontaneous Ca^2+^ release under stress conditions, suggesting that JCN may function as a brake stabilizing RyR2 activity^22^. In failing hearts, JCN protein is decreased^23^. Decreased JCN may increase RyR2 activity and Ca^2+^ release from sarco/endoplasmic reticulum, which promotes cardiac injury^22^. In fact, a previous study reported that ablation of JCN increased I/R injury through induction of endoplasmic reticulum stress and calpain activation in the heart^24^. As both JCN and JPH2 share similar functions in the regulation of RyR2 activity, it is intriguing to find out if JPH2 regulates RyR2-dependent Ca^2+^ release through JCN in cardiomyocytes.

This study determined if JPH2 plays a role in cardiomyocyte survival/death and explored the mechanisms by which JPH2 modulates JCN in maintaining cytosolic Ca^2+^ homeostasis. We report that JPH2 is required for the stability of JCN and that a reduction of JPH2 predisposes JCN to E3 ubiquitin ligase muscle RING-finger protein-1 (Murf1)-mediated ubiquitination and proteasome-dependent degradation, leading to increased cytosolic Ca^2+^ and cardiomyocyte death. Accordingly, over-expression of JPH2 and JCN protects cardiomyocytes against stress-induced death. These findings demonstrate for the first time that both JPH2 and JCN are survival factors of cardiomyocytes and provide molecular basis for JPH2 stabilization of JCN in maintaining cytosolic Ca^2+^ homeostasis.

## Methods

### Animals

This research conformed to the Guide for the Care and Use of Laboratory Animals published by the US National Institute of Health (NIH Publication, 8^th^ Edition, 2011). All experimental protocols were approved by the Animal Use Subcommittee at Soochow University, China, and Western University, Canada. Breeding pairs of C57BL/6 mice were purchased from the Cyagen Biosciences (Suzhou, China) and the Jackson Laboratory (Bar Harbor, ME, USA) to produce neonates for cardiomyocyte isolation. Neonatal rats born within 48 hours were purchased from Shanghai Lab-Animal Research Center, Shanghai, China. All animals were housed in a temperature- and humidity-controlled facility on a 12:12 h light: dark cycle with water and food *ad libitum*.

### Cell culture, adenoviral infection and DNA transfection

The neonatal cardiomyocytes were prepared from mice or rats born within 24 or 48 hours, respectively after decapitation and cultured according to methods we previously described^25^. Adult mice were euthanized by cervical dislocation and their cardiomyocytes were isolated and cultured as we previously described^26^. The rat cardiomyocyte-like H9c2 cells were purchased from the American Type Culture Collection (ATCC) and cultured H9c2 cells were employed within 10 generations.

Cardiomyocytes were infected with an adenoviral vector containing human *JPH2* (Ad-JPH2, Vector Biolabs, Malvern, PA, USA), shRNA for mouse *JPH2* (Ad-shJPH2, SignaGen, Rockville, MD, USA), mouse *JCN* (Ad-JCN), shRNA for rat Murf1 (Ad-shMurf1, Hanbio Biotechnology Co., Ltd., Shanghai, China) or antisense against mouse *JCN* (Ad-asJCN) at a multiplicity of infection of 100 PFU/cell. An adenoviral vector containing green fluorescent protein (Ad-GFP, SignaGen, Rockville, MD, USA) or hemagglutinin (Ad-HA, Vector Biolabs, Malvern, PA, USA) served as controls as appropriate. Adenovirus-mediated gene transfer was performed as previously described ^25^.

H9c2 cells were transfected with plasmid containing DDK-tagged mouse *JCN* (Accession number: NM_133723, Origene, Rockville, MD, USA) or its mutant (K8R/K102R/K107R/K140R, generated by Norclone Biotech Laboratories, London, ON. Canada) using the jetPRIME DNA transfection reagent (Polyplus-Transfection, Illkirch, France) according to the manufacturer’s instructions. A plasmid containing EGFP (*p*CMV-EGFP) served as a control. Hypoxia/re-oxygenation on H9c2 cells was conducted as previously described ^27^.

For incubation with oleate and palmitate, oleate and palmitate were dissolved in distilled water and then conjugated with bovine serum album (Sigma-Aldrich Canada Co., Oakville, ON, Canada). Neonatal cardiomyocytes, adult cardiomyocytes and H9c2 cells were exposed to culture medium with oleate and palmitate at concentrations of 0.8 mM (the fatty acid to album is about 1:5), 0.2 mM, and 0.4 mM, respectively. These doses for oleate and palmitate were based on our preliminary study.

The dose of doxorubicin (1 μM) was chosen based on our recent report ^28^.

### Bimolecular fluorescence complementation (BiFC) assay

A BiFC assay was peroformed to determine the interaction between JPH2 and JCN protein ^29^. Plasmids pVN155 and pVC155 were purchased from Addgene (Watertown, MA, USA). The coding regions of human *Jph2* cDNA (accession number: NM_020433, sharing 91% homology with mouse JPH2 protein) and mouse *Jcn* cDNA (Accession number: NM_133723, Origene, Rockville, MD, USA) were inserted into pVN155 and pVC155, respectively. The resulting plasmids (pVN155/*Jph2* and pVC155/*Jcn*) were co-transfected into H9c2 cells using the jetPRIME DNA transfection reagent (Polyplus-Transfection, Illkirch, France) according to the manufacturer’s instructions. Plasmids pVN155+pVC155, pVN155+pVC155/*Jcn*, or pVN155/*Jph2*+pVC155 were co-transfected into H9c2 cells as controls, respectively. Twenty-four hours after transfection, the cells were monitored for GFP signal under fluorescence microscope and nuclei were stained blue by Hoechst 33342 (Thermo Fisher Scientific, Mississauga, ON, Canada).

### Co-Immunoprecipitation

Co-Immunoprecipitation analyzed protein-protein interactions. Briefly, JPH2 or JCN and their interacting proteins were co-precipitated by using an immunoprecipitation kit (Dynabeads Protein G; Life Technologies, Inc., ON, Canada) in tissue or cell lysates according to the manufacturer’s instructions. Both JPH2 and JCN interacting proteins were subjected to western blot analysis and/or mass spectrometric analysis.

### Mass spectrometry

JPH2 or JCN and their interacting proteins were co-precipitated and then subjected to mass spectrometry.Briefly, fractions containing tryptic peptides were resolved and ionized using nanoflow UHPLC (Easy-nLC 1200 system, Thermo Fisher Scientific Inc., USA) coupled to an Orbitrap Fusion Lumos Tribrid mass spectrometer (Thermo Fisher Scientific Inc., USA). After nanoflow chromatography and electrospray ionization using an Acclaim PepMap 100 C18 HPLC column (100 Å, 2μm, 50 μm id × 150 mm, Thermo Fisher Scientific Inc., USA), peptide mixtures were injected onto the column at a flow rate of 300 nL/min using linear gradients from 1 to 100% v/v aqueous acetonitrile in 0.1% v/v formic acid over 60 minutes. The data dependent mode of mass spectrometer was cycle time (time between master scans: 3 secs), with a resolution of 60,000 and m/z range of 350–1600. Data were processed using Proteome Discoverer 1.2 (Thermo Fisher Scientific Inc., USA), and relevant targeted proteins were searched using SEQUEST (Thermo Fisher Scientific Inc., USA) in a Uniprot (uniprot.org) mouse database. Search parameters included a precursor mass tolerance of 10 ppm and a fragment mass tolerance of 0.02 Dalton.

The ubiquitin modifications of JCN were identified by mass spectrometry through commercial service provided by Applied Protein Technology Inc. (Shanghai, China).

### Western blot analysis

Heart tissue lysates or cell lysates were subjected to SDS-polyacrylamide gel electrophoresis. After the proteins were transferred to PVDF membranes, western blot analysis determined the protein levels of interest using specific antibodies. Primary antibodies against JPH2 (catalog number: ab79071, 1:1000 dilution, Abcam, Cambridge, UK), JCN (catalog number: PA5-43688, 1:1000 dilution, Invitrogen Canada Inc., Burlington, ON, Canada), DDK (catalog number: PA1-984B, 1:1000 dilution, Invitrogen Canada Inc., Burlington, ON, Canada), LC3B (catalog number: 2775S, 1:1000 dilution, Cell Signaling Technology, Danvers, MA, USA), MURF1 (catalog number: ab172479, 1:1000 dilution, Abcam, Cambridge, UK), ubiquitin (catalog number: 3933S, 1:1000 dilution, Cell Signaling Technology, Danvers, MA, USA), and GAPDH (catalog number: 2118S, 1:1000 dilution, Cell Signaling Technology, Danvers, MA, USA) were employed as previously described ^25^. Horseradish peroxidase (HRP)-conjugated goat anti-mouse IgG (H+L) secondary antibody was purchased from Bio-Rad (catalog number: 1706516, Hercules, CA, USA), and HRP-conjugated goat anti-rabbit IgG (H+L) was purchased from Bio-Rad (catalog number: 1706515, Hercules, CA, USA).

### Active caspase-3

As described previously ^30^, caspase-3 activity in cell lysates was measured using a Caspase-3 Fluorescence Assay Kit (Biomol Research Laboratories, Inc., Plymouth, PA, USA).

### Determination of cellular DNA fragmentation

Cultured H9c2 cells were pre-labeled with BrdU (1 μg/mL). DNA fragmentation was evaluated using a Cellular DNA Fragmentation ELISA Kit (Roche Applied Science, Mannheim, Germany) according to the manufacturer’s instructions.

### LDH release

Lactate dehydrogenase (LDH) activity in culture medium was measured using a commercially available LDH Cytotoxicity Assay Kit (Cayman Chemical, Ann Arbor, MI, USA) following the manufacturer’s instructions.

### Annexin V staining

Cell death was determined in adult mouse cardiomyocytes by FITC-conjugated annexin V staining as described previously ^31^ (Invitrogen Canada Inc., Burlington, ON, Canada). Nuclei were stained by Hoechst 33324 (Invitrogen Canada Inc., Burlington, ON, Canada).

### Calpain activity

Calpain activities were measured in CMVEC lysates using a fluorescence substrate *N*-succinyl-LLVY-AMC (Cedarlane Laboratories, Burlington, ON, Canada) as previously described ^32^.

### Cytosolic Ca^2+^ measurement

Cytosolic free Ca^2+^ was determined by the cell-permeable Ca^2+^ imaging dye Fura-2 AM (eBioscience, CA, USA)^33^. The fluorescence intensity was measured by a fluorescence spectrophotometer with excitation of 340/380 nm and emission of 510 nm. Finally, the concentration of cytosolic free Ca^2+^ was calculated according to the formula: [Ca^2+^]_i_ = Kd ×[(R-R_EGTA_)/(R_Triton-X 100_ – R)] × (F380 _EGTA_/F380 _Triton-X 100_). (Kd = 225 nM, R = F340/F380)

### Real-time reverse transcription PCR

Total RNA was extracted using TRIzol Reagent (Sigma-Aldrich Canada Co., Oakville, ON, Canada) following the manufacturer’s instructions. Real-time reverse transcription PCR analyzed the mRNA levels of JCN and GAPDH as previously described ^34^. The sequences of the primers for the *Jph2*, *Jcn* and *Gapdh w*ere as follows. *Jph2*: 5’-ACCCAGGAGGATGAGAGGTT-3’ (forward), 5’-CAGCTGGGCTTCTTTGTTTC-3’ (reverse); *Jcn*: 5’-GTCGTGTGGTTTGACTTGGT-3’ (forward), 5’-ATGCTTTCTGGGATGTGTCTGA-3’ (reverse); and *Gapdh*: 5’-CAGACTCTGCGATGTTTCCA-3’ (forward), 5’-GCCTGAGCACTTCCAGAAAC-3’ (reverse).

### Statistical analysis

The data are expressed as the mean ± SD. Student’s *t* test was used to compare the data between two groups. ANOVA followed by the Newman-Keuls test was used for multi-group comparisons. A *P* value of less than 0.05 was considered significant.

## Results

### Conditions of stress reduce JPH2 expression through aberrant autophagy

To determine if conditions of stress decreased JPH2 expression in cardiomyocytes, we exposed neonatal mouse cardiomyocytes to palmitate (800 μM) or doxorubicin (1 μM). Western blot analysis showed that exposure to palmitate or doxorubicin resulted in lower levels of JPH2 protein (**Figures A** and **B)**, which were correlated with a reduction of its mRNA level (**Figures C** and **D)**. Thus, different conditions of stress reduce JPH2 expression in cardiomyocytes at the mRNA levels.

Since our recent study found that aberrant autophagy mediated a reduction of JPH2 protein in calpain-2 over-expressing cardiomyocytes ^35^, we determined the alterations of autophagy in cardiomyocytes. We showed that palmitate induced autophagic flux blockage in cardiomyocytes (**Figure E**). Our previous study also reported autophagic flux blockage in cardiomyocytes in response to doxorubicin ^28^. Notably, inhibition of autophagy with 3-MA (500 μM) attenuated the reduction of *Jph2* mRNA levels in palmitate or doxorubicin-stimulated cardiomyocytes (**Figures C** and **D**), suggesting that aberrant autophagy may be an important mechanism contributing to down-regulation of JPH2 expression in cardiomyocytes under stress.

### JPH2 over-expression prevents apoptosis in cardiomyocytes under stress

To define the role of JPH2 in cardiomyocyte survival/death under stress, we infected neonatal mouse cardiomyocytes with Ad-JPH2 or Ad-HA as a control. Twenty-four hours later, cardiomyocyte death was induced by incubation with palmitate (800 μM) or doxorubicin (1 μM) for 24 hours. Oleate and saline served as controls for palmitate and doxorubicin, respectively. Infection with Ad-JPH2 resulted in higher JPH2 protein levels in cardiomyocytes (**Figure 2A**). Both palmitate and doxorubicin induced greater caspase-3 activity and DNA fragmentation in Ad-HA-infected cardiomyocytes compared with oleate and saline, respectively; however, their levels were much lower in Ad-JPH2-infected cardiomyocytes after exposure to palmitate and doxorubicin, indicating an inhibitory effect of JPH2 over-expression on palmitate or doxorubicin-induced apoptosis (**Figures B-E**). The inhibitory effect of JPH2 over-expression on palmitate-induced cell death was recapitulated in adult mouse cardiomyocytes (**Figure 2F** and **2G**) and neonatal rat cardiomyocytes (**Figure S1A** and **S1B**), where palmitate resulted in lower protein levels of JPH2 (**Figure S1C**). Similarly, in an *in vitro* model of hypoxia/re-oxygenation in H9c2 cells, we showed a similar inhibitory effect of JPH2 over-expression on caspase-3 activity and DNA fragmentation (**Figure S2)**. These findings indicate that JPH2 prevents cardiomyocyte death under stress.

**Figure 1.**
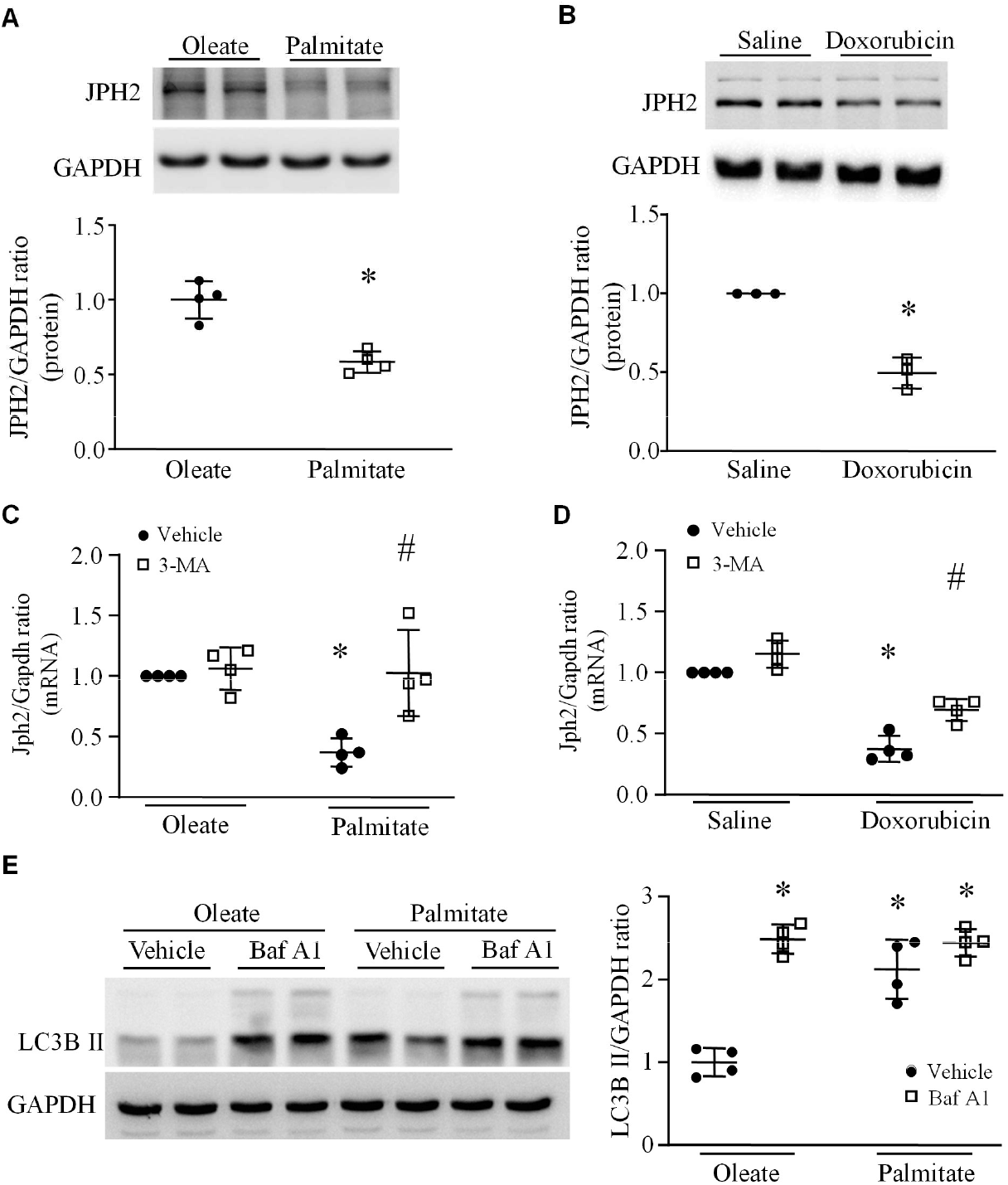
JPH2 expression and autophagic flux analysis. Neonatal cardiomyocytes were incubated with palmitate or oleate, or doxorubicin or saline for 24 hours. The levels of JPH2 protein and mRNA were determined by western blot analysis and real-time RT-PCR. (**A** and **B**) the upper panel is representative western blots for JPH2 and GAPDH and lower panel is quantification of JPH2/GAPDH ratio. (**C** and **D**) the mRNA levels of *Jph2* relative to *Gapdh*. (**E**) After palmitate incubation for 24 hours, cardiomyocytes were incubated with bafilomycin A1 or vehicle for 2 hours. The protein levels of LC3BII and GAPDH were determined by western blot analysis. Left panel: representative western blots for LC3BII and GAPDH; right panel: quantification of LC3BII/GAPDH ratio. Data are mean ± SD, n = 3-4. \**P*< 0.05 versus oleate, oleate + vehicle or saline + vehicle, and #*P*< 0.05 versus palmitate + vehicle or doxorubicin + vehicle.

**Figure 2.**
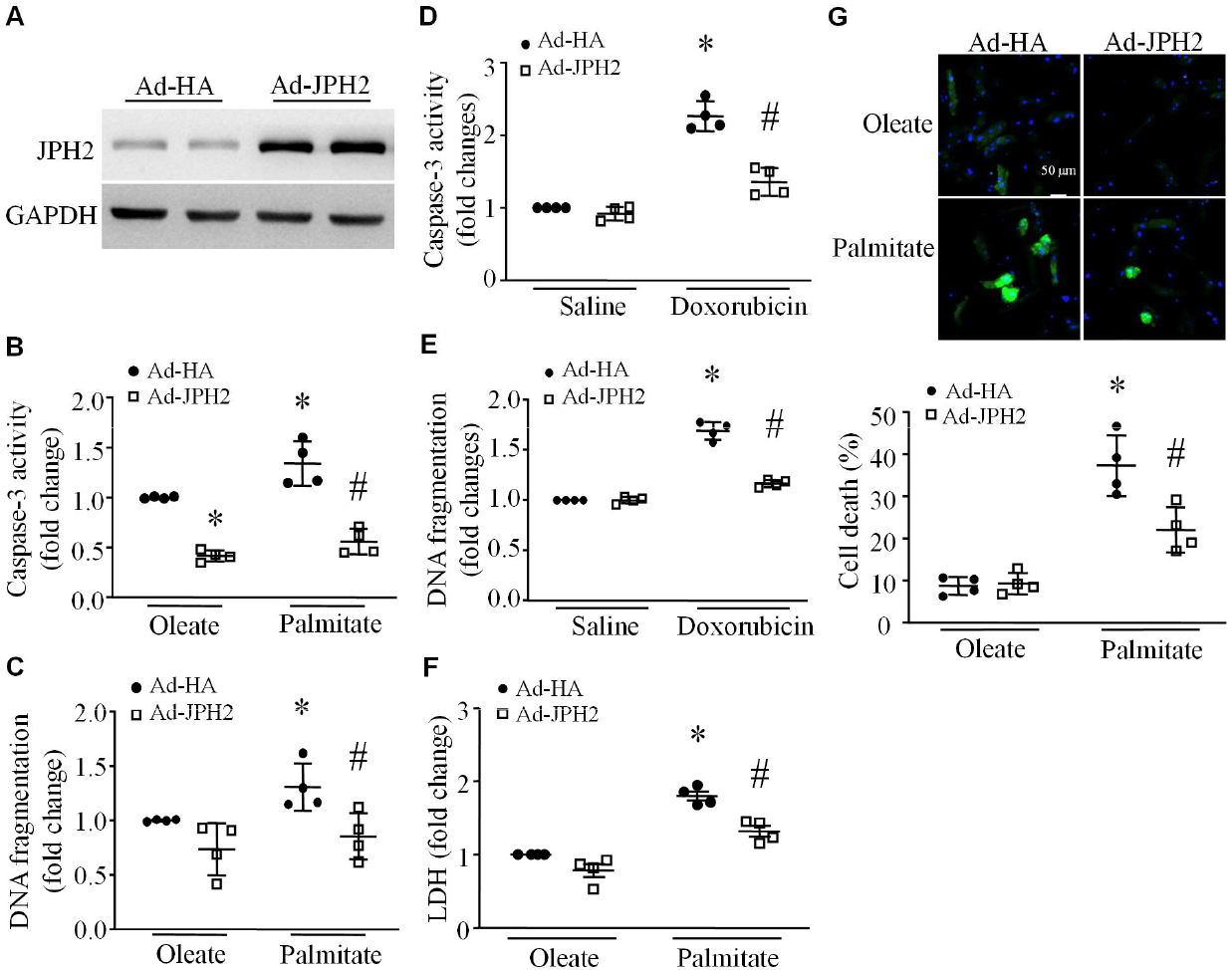
Assessment of cell death. Neonatal and adult mouse cardiomyocytes were infected with Ad-JPH2 or Ad-HA, and then exposed to palmitate or oleate, or doxorubicin or saline for 24 hours. (**A**) A representative western blot for JPH2 protein expression in neonatal mouse cardiomyocytes from 3 different cultures. (**B** and **C**) Apoptosis induced by palmitate in neonatal mouse cardiomyocytes was assessed as caspase-3 activity (**B**) and DNA fragmentation (**C**). (**D** and **E**) Apoptosis induced by doxorubicin in adult cardiomyocytes was assessed as caspase-3 activity (**D**) and DNA fragmentation (**E**). (**F** and **G**) Effect of JPH2 over-expression on palmitate-induced cell death in adult cardiomyocytes. (**F**) LDH release. (**G**) Upper panel: representative pictures for annexin V staining (green color for annexin V and blue color for nuclei) and bottom panel: quantification of annexin V staining positive cells. Data are mean ± SD, n = 4. \**P*< 0.05 versus oleate + Ad-HA or saline + Ad-HA and #*P*< 0.05 versus palmitate + Ad-HA or doxorubicin + Ad-HA.

### Over-expression of JPH2 prevents aberrant cytosolic Ca^2+^ level in cardiomyocytes under stress

Disturbance of cytosolic Ca^2+^ has been implicated in cardiomyocyte death ^36^. JPH2 was reported to stabilize T-tubule organization and RyR2 calcium channel in cardiomyocytes and decreased JPH2 leads to excitation-contraction un-coupling and Ca^2+^ leakage via RyR2^4–6, 8, 9, 11, 13, 19, 20, 37^. We then analyzed cytosolic Ca^2+^ levels in cardiomyocytes under stress. The level of cytosolic Ca^2+^ was much higher in palmitate- or doxorubicin-compared with oleate- or saline-stimulated neonatal mouse cardiomyocytes, respectively (**Figures 3A** and **3B)**. However, over-expression of JPH2 by infection of Ad-JPH2 resulted in a lower level of cytosolic Ca^2+^ in neonatal cardiomyocytes compared to Ad-HA infection. In adult mouse cardiomyocytes, palmitate-induced increase in cytosolic Ca^2+^ levels was also attenuated by JPH2 over-expression **(Figure 3C**).

**Figure 3.**
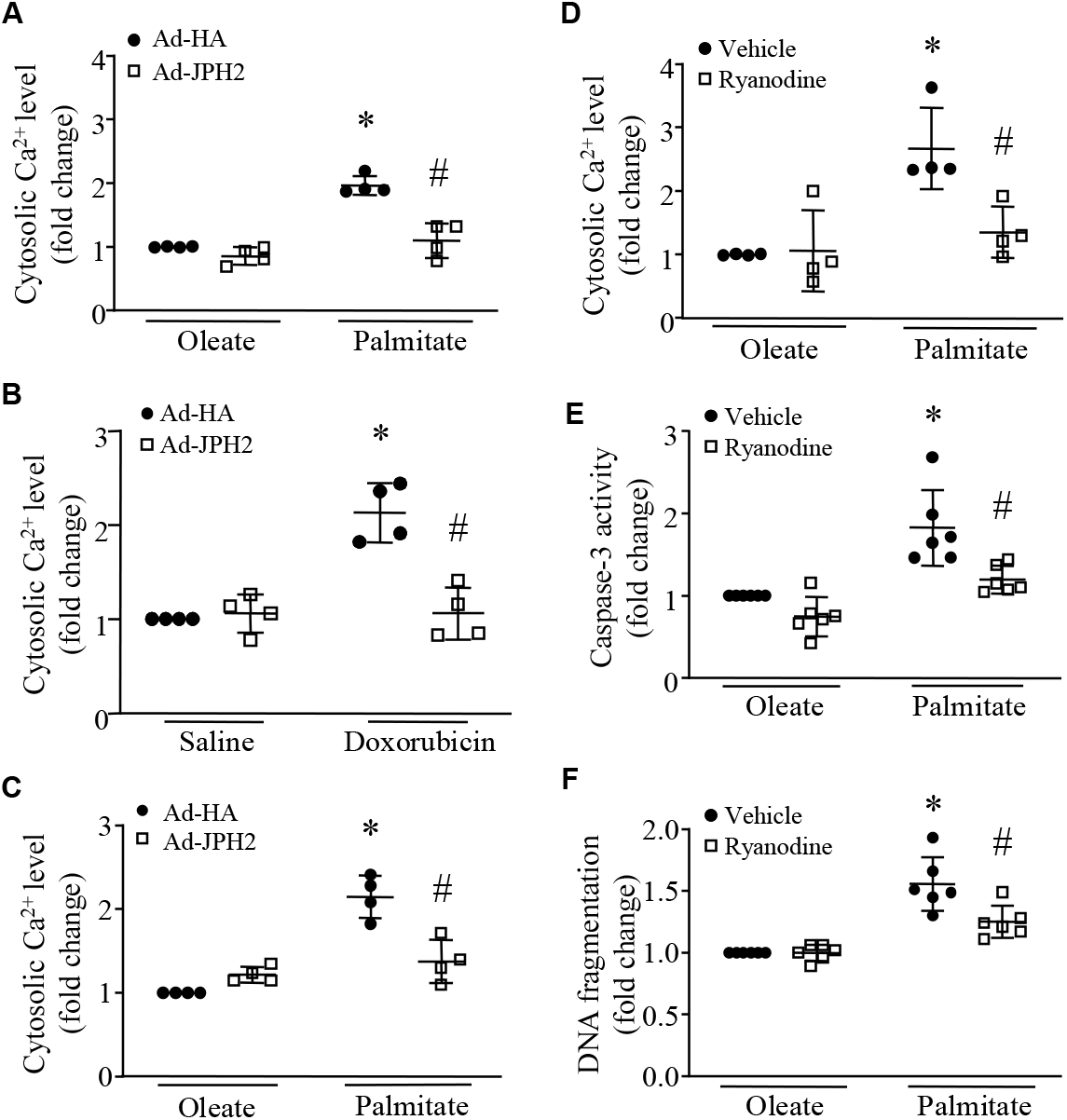
Role of JPH2 in the regulation of cytosolic Ca^2+^ in cardiomyocytes. (**A**–**C**) Neonatal or adult mouse cardiomyocytes were infected with Ad-JPH2 or Ad-HA and then exposed to palmitate or oleate, or doxorubicin or saline for 24 hours. Cytosolic Ca^2+^ in neonatal cardiomyocytes (**A** and **B**) and adult cardiomyocytes (**C**) were analyzed. (D**-F**) Neonatal mouse cardiomyocytes were incubated with palmitate or oleate in the presence of Ryanodine (50 μM) or Vehicle for 24 hours. Cytosolic Ca^2+^ (**D**), caspase-3 (**E**) and DNA fragmentation were determined (**F**). Data are mean ± SD, n = 4-6 independent cell cultures. \**P*< 0.05 versus oleate + Ad-HA, oleate + vehicle, or saline + Ad-HA and #*P*< 0.05 versus palmitate + Ad-HA, palmitate + vehicle, or doxorubicin + Ad-HA.

We then determined whether inhibition of RyR2 reduced cytosolic Ca^2+^ levels thereby preventing apoptosis in stressed cardiomyocytes. To address this, we incubated neonatal mouse cardiomyocytes with palmitate or oleate in combination with vehicle or Ryanodine (50 μM), a selective inhibitor of RyR2 (Note: low concentration of ryanodine at nM range activates RyR2). Ryanodine attenuated cytosolic Ca^2+^ level in palmitate-induced cardiomyocytes (**Figure 3D**). Ryanodine concomitantly reduced apoptosis induced by palmitate in cardiomyocytes (**Figure 3E** and **3F**). Taken together, these results indicate that JPH2 prevents cardiomyocyte apoptosis at least partially by reducing RyR2-dependent Ca^2+^ release.

### Knockdown of JPH2 sufficiently increases cytosolic Ca^2+^ and induces apoptosis in cardiomyocytes

Next, we examined if decreased JPH2 was sufficient to induce apoptosis. Accordingly, we knocked down JPH2 in neonatal mouse cardiomyocytes by infection of an adenoviral vector containing shRNA for *JPH2* (Ad-shJPH2). Infection of Ad-shJPH2 reduced JPH2 protein levels (**Figure 4A**), and concomitantly increased caspase-3 activity and DNA fragmentation in cardiomyocytes (**Figure 4B** and **4C**). Similar to conditions of stress, Ad-shJPH2-mediated knockdown of JPH2 was sufficient to elevate cytosolic Ca^2+^ level in neonatal mouse cardiomyocytes (**Figure 4D**).

**Figure 4.**
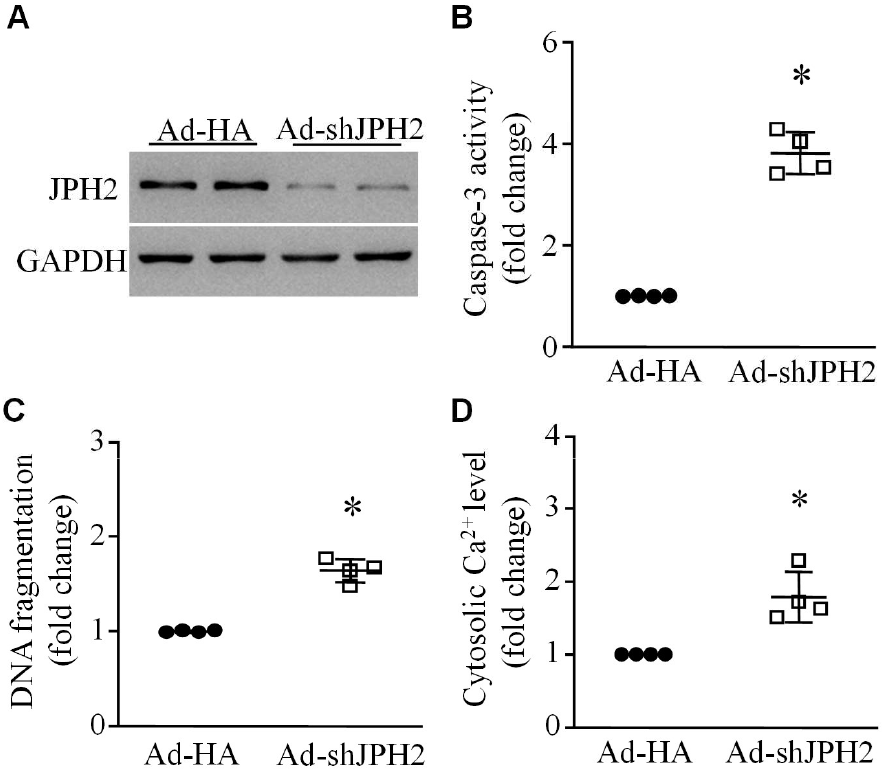
Effects of JPH2 knockdown on cytosolic Ca^2+^ and apoptosis. Neonatal mouse cardiomyocytes were infected with Ad-shJPH2 or Ad-HA. (**A**) A representative western blot for JPH2 protein expression from 3 different cultures. (**B**) Caspase-3 activity. (**C**) DNA fragmentation. (**D**) Cytosolic Ca^2+^ levels. Data are mean ± SD, n = 4 independent cell cultures. \**P*< 0.05 versus Ad-HA.

### JPH2 is required for JCN protein stability in cardiomyocytes

Studying the downstream mechanisms by which JPH2 attenuated cytosolic Ca^2+^ levels and apoptosis in cardiomyocytes, we analyzed JPH2 immunoprecipitates from mouse heart tissue lysates by mass spectrometry and identified JCN as one JPH2 interacting protein (**Figure 5A1**). Likewise, JPH2 was present in JCN immunoprecipitates (**Figure 5A2**). Immunoprecipitation followed by western blot analysis verified the physical interaction between JPH2 and JCN (**Figure 5B**), which was further confirmed by the bimolecular fluorescence complementation (BiFC) assay (**Figure 5C1-3**). Notably, both palmitate and JPH2 knockdown resulted in lower protein levels of JCN in neonatal mouse cardiomyocytes without changing its mRNA levels (**Figure 6A-6D**). Furthermore, over-expression of JPH2 by Ad-JPH2 preserved the protein levels of JCN in neonatal mouse cardiomyocytes under stress (**Figure 5E**). These results suggest that conditions of stress may accelerate JCN protein degradation and JPH2 may be important in maintaining JCN protein stability. Indeed, cycloheximide pulse chase assay revealed that palmitate significantly shortened the half-life of JCN protein (**Figure 5F**), which was mitigated by JPH2 over-expression in neonatal mouse cardiomyocytes (**Figure 5G**).

**Figure 5.**
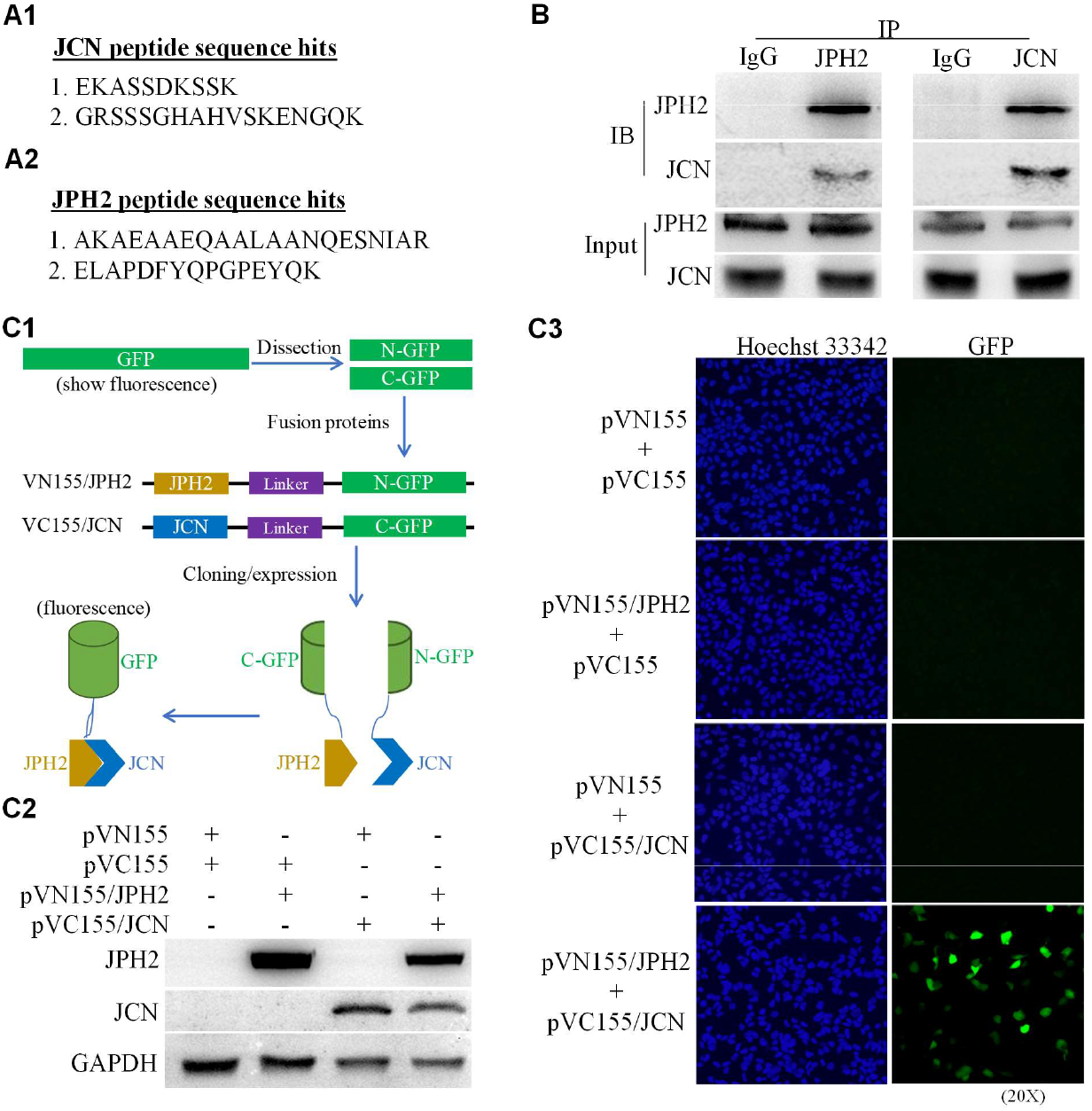
JPH2 and JCN interactions. (**A**) Identification of JCN peptide sequences in JPH2 immunoprecipitates (**A1**) and JPH2 peptide sequences in JCN immunoprecipitates from mouse hearts tissue lysates by mass spectrometry (**A2**). (**B**) Mouse heart tissue lysates were subjected to immunoprecipitation using anti-JPH2 antibody or anti-JCN antibody followed by western blot analysis using anti-JPH2 antibody or anti-JCN antibody. IgG served as a negative control for immunoprecipitation. Representative western blots for JPH2 and JCN are presented. (**C**) Bimolecular fluorescence complementation (BiFC) assay. (**C1**) Structures of plasmids *p*VN155/JPH2 and *p*VC155/JCN for the BiFC assay. (**C2**) A representative western blot for JPH2 and JCN expression in H9c2 cells after transfection with plasmid *p*VN155/JPH2 and *p*VC155/JCN, respectively. (**C3**) A representative GFP fluorescence micro-photograph in H9c2 cells after co-transfection with *p*VN155 and *p*VC155, *p*VN155/JPH2 and *p*VC155, *p*VN155 and *p*VC155/JCN, or *p*VN155/JPH2 and *p*VC155/JCN, respectively. The experiments in B, C2 and C3 were repeated 3 times using 3 different cell cultures.

**Figure 6.**
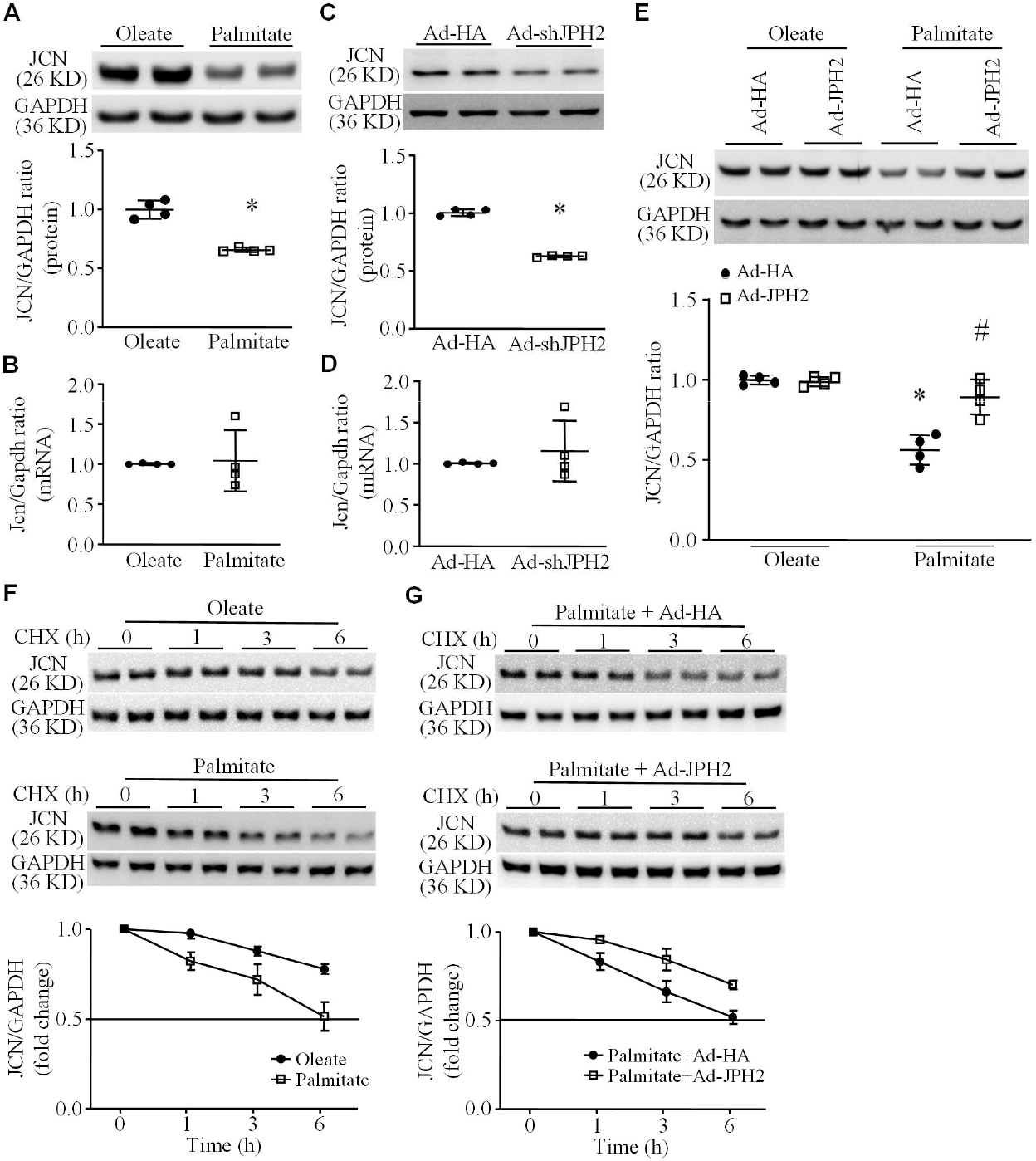
JPH2 regulation of JCN stability. (**A-D**) Neonatal mouse cardiomyocytes were incubated with palmitate or oleate, or infected with Ad-shJPH2 or Ad-HA for 24 hours. The levels of JCN protein (**A** and **C**) and mRNA were analyzed (**B** and **D**). (**E**) Neonatal mouse cardiomyocytes were infected with Ad-JPH2 or Ad-HA and then exposed to palmitate or oleate for 24 hours. Upper panel: A representative western blot for JCN, and bottom panel: quantification of JCN protein levels relative to GAPDH. Data are mean ± SD, n = 4 different cell cultures. \**P*< 0.05 versus oleate, Ad-HA or oleate + Ad-HA and #*P*< 0.05 versus oleate + Ad-HA. (**F** and **G**) Analysis of JCN protein stability by cycloheximide pulse chase assay. Neonatal mouse cardiomyocytes were infected with Ad-JPH2 or Ad-HA and then exposed to palmitate or oleate. Sixteen hours later, cardiomyocytes were incubated with cycloheximide for different time periods and the protein levels of JCN were determined by western blot analysis. (**F**) Effect of palmitate on JCN protein half-life. (**G**) Effect of JPH2 over-expression on JCN protein half-life in palmitate-incubated cardiomyocytes. Upper panel: Representative western blots for JCN protein. Bottom panel: Quantification of JCN proteins. Data are mean ± SD, n = 4 different cell cultures.

### JPH2 prevents JCN ubiquitination and proteasome-dependent degradation in cardiomyocytes under stress

Since the ubiquitin-proteasome system accounts for the degradation of majority of cellular proteins^38^, we determined whether the proteasomal degradation pathway contributed to the reduction of JCN protein level. We incubated neonatal mouse cardiomyocytes with palmitate or oleate for 24 hours in the presence of vehicle or MG132 (10 μM), a selective inhibitor of proteasomes. As shown in **Figure 7A**, Co-incubation with MG132 relatively preserved the levels of JCN protein in palmitate-induced cardiomyocytes, suggesting the involvement of the proteasomal degradation pathway. We then determined the states of JCN ubiquitination in this context. To do this, we over-expressed DDK-tagged JCN and JPH2 in H9c2 cells by transfection of a plasmid expressing DDK-tagged JCN and infection of Ad-JPH2, respectively, and then incubated the cells with palmitate or oleate. We chose H9c2 cells because it is extremely hard to transfect a plasmid DNA into primary cardiomyocytes. Palmitate induced more ubiquitination of DDK-tagged JCN protein, which was prevented by over-expression of JPH2 (**Figure 7B**).

**Figure 7.**
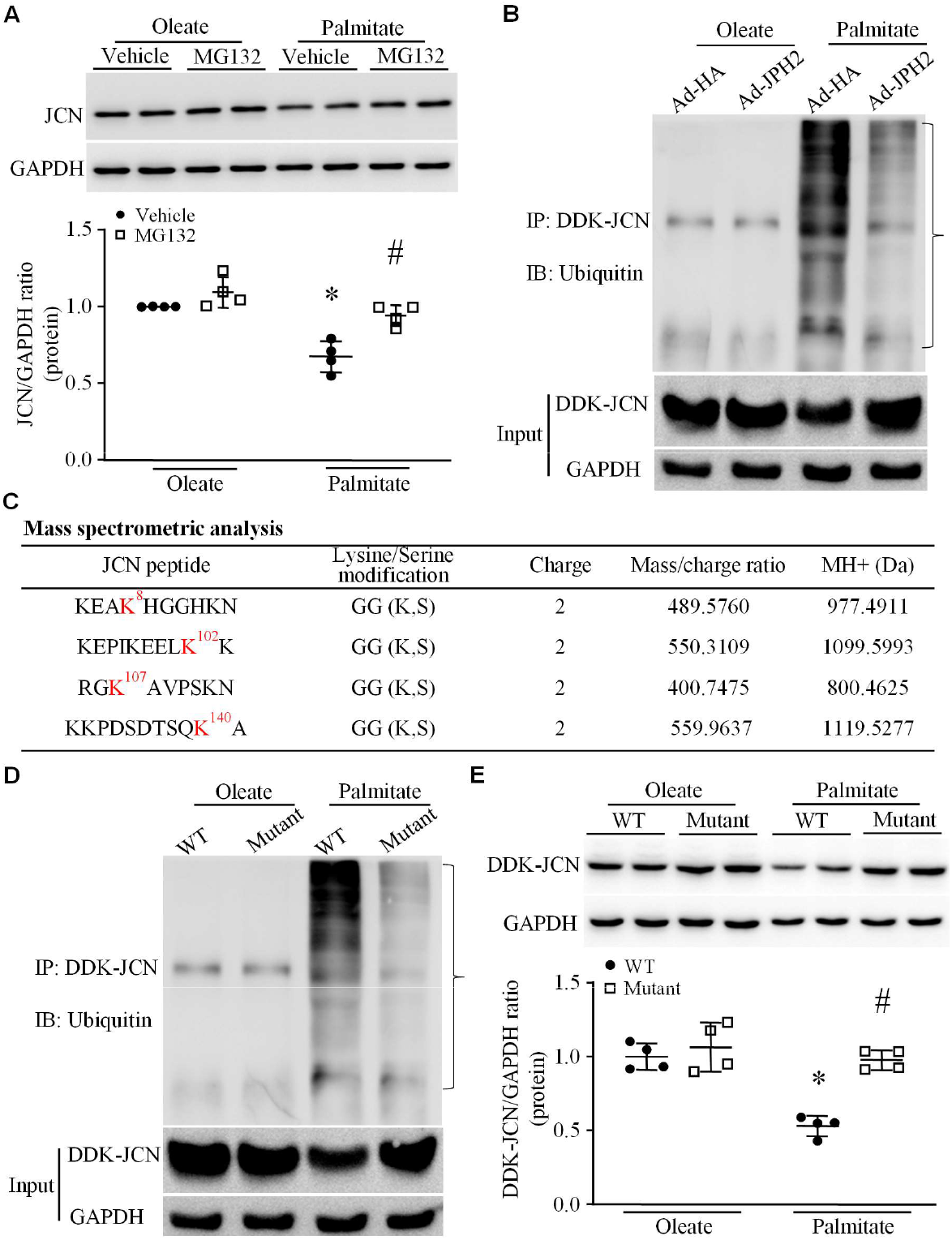
Effects of JPH2 over-expression on palmitate-induced JCN ubiquitination and proteasomal degradation. (**A**) Neonatal mouse cardiomyocytes were incubated with palmitate or oleate in the presence of MG132 or vehicle for 24 hours. Upper panel: a representative western blot for JCN. Bottom panel: quantification of JCN protein levels relative to GAPDH. (**B**) H9c2 cells were transfected with plasmid expressing DDK-tagged JCN and then infected with Ad-JPH2 or Ad-HA. After incubation with palmitate or oleate for 24 hours, DDK-JCN protein was pulled down by immunoprecipitation using anti-DDK antibody, and western blot analysis determined DDK-JCN ubiquitination using anti-ubiquitin antibody. A representative western blot for the ubiquitinated DDK-JCN protein from one out of 4 different cell cultures is presented. (**C**) Mass spectrometry identified JCN protein ubiquitination sites in palmitate-induced neonatal rat cardiomyocytes. (**D** and **E**) H9c2 cells were transfected with plasmids expressing wild-type DDK-tagged JCN (WT) or a mutant of DDK-tagged JCN (K8A/K102A/K107A/K140A, mutant), and then exposed to palmitate or oleate for 24 hours. (**D**) DDK-JCN protein was pulled down by immunoprecipitation using anti-DDK antibody, and its ubiquitination was then determined by western blot analysis using anti-ubiquitin antibody. A representative western blot for the ubiquinated DDK-JCN protein from one out of 4 different cell cultures is presented. (**E**) Upper panel: representative western blot for DDK-JCN protein and bottom panel: quantification of DDK-JCN protein levels relative to GAPDH. Data are mean ± SD, n = 4 different cell cultures. \**P*< 0.05 versus oleate + vehicle or oleate + WT and #*P*< 0.05 versus palmitate + WT or palmitate + vehicle.

To provide further insights into ubiquitin-mediated proteasomal degradation of JCN, our in-silico analysis of JCN protein with Bayesian Discriminant Method identified multiple potential ubiquitination sites (**Figure S3**), some of which were verified by analyzing JCN immunoprecipitates from palmitate-induced neonatal cardiomyocytes using mass spectrometry (**Figure 7C**). To determine putative ubiquitination sites which were modulated under stress, we generated a mutant of DDK-tagged JCN (muJCN) by replacing lysine with arginine at 4 different ubiquitination sites (K8A, K102A, K107A and K140A) on JCN protein, which were identified by in-silico analysis and observed in palmitate but not oleate-induced cardiomyocytes. After transfection of plasmids expressing JCN or muJCN, both of which were DDK-tagged fusion proteins, we incubated H9c2 cells with palmitate or oleate for 24 hours. Immunoprecipitation followed by western blot analysis showed that palmitate induced dramatically higher ubiquitination of DDK-tagged JCN but not of muJCN (**Figure 7D**). Similarly, palmitate lowered DDK-tagged JCN protein levels in H9c2 cells; however, the protein levels of DDK-tagged muJCN were relatively preserved in palmitate-incubated H9c2 cells (**Figure 7E**). These results provide direct evidence demonstrating that JPH2 prevents ubiquitin-mediated proteasomal degradation of JCN in cardiomyocytes under stress.

### JPH2 blocks Murf1-mediated JCN ubiquitination and degradation in cardiomyocytes

To explore the mechanisms of JCN ubiquitination, we incubated neonatal rat cardiomyocytes with palmitate for 24 hours and analyzed JCN interacting proteins from its immunoprecipitates of cell lysates by mass spectrometry. An E3 ubiquitin ligase, Murf1, was found in JCN immunoprecipitates (**Figure 8A**). To examine the role of Murf1, we infected neonatal rat cardiomyocytes with an adenoviral vector containing shRNA for rat Murf1 or Ad-HA as a control and then incubated them with palmitate or oleate. Knockdown of Murf1 resulted in higher protein levels of JCN (**Figure 8B** and **Figure S4**) and prevented JCN ubiquitination in palmitate-overloaded cardiomyocytes (**Figure 8C**). Similarly, incubation with pharmacological inhibitor of Murf1 EMBL (10 μM) normalized the protein levels of JCN in palmitate-stimulated cardiomyocytes (**Figure 8D**). Furthermore, as indicated by the cross-pulldown assay, incubation with palmitate increased Murf1-JCN interaction in neonatal mouse cardiomyocytes, which was prevented by JPH2 over-expression (**Figure 8E**). To further address the role of JPH2 in this context, knockdown of JPH2 resulted in an increase in Murf1-JCN interaction in cardiomyocytes (**Figure 8F**). Thus, we speculate that intact JPH2-JCN relationship prevents Murf1-JCN interaction thereby disabling Murf1-mediated JCN ubiquitination in cardiomyocytes under stress.

**Figure 8.**
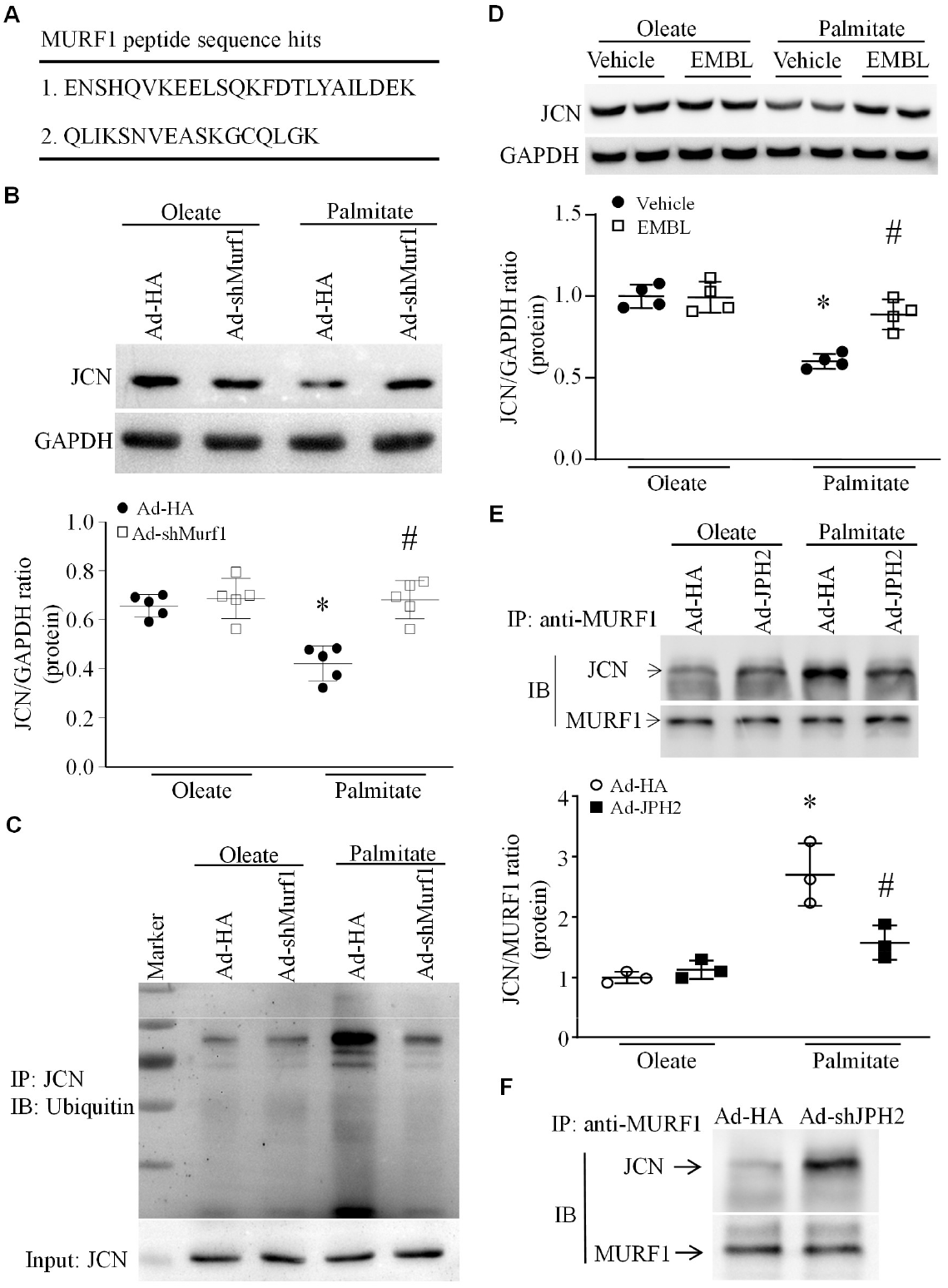
Role of Murf1 in JCN ubiquitination and degradation. (**A**) Neonatal rat cardiomyocytes were incubated with palmitate for 24 hours. Immunoprecipitation was performed to pull down JCN and its interacting proteins, which were analyzed by mass spectrometry. Two MURF1 peptide sequence hits were identified in JCN immunoprecipitates. (**B** and **C**) Neonatal rat cardiomyocytes were infected with an adenoviral vector containing shRAN for rat Murf1 (Ad-shMurf1) or Ad-HA and then exposed to palmitate or oleate for 24 hours. (**B**) Upper panel: a representative western blot for JCN and bottom panel: quantification of JCN protein levels relative to GAPDH. Data are mean ± SD, n = 5 different cell cultures. * *P*< 0.05 versus oleate + Ad-HA and # *P*< 0.05 versus palmitate + Ad-HA. (**C**) JCN protein was pulled down by immunoprecipitation using anti-JCN antibody, followed by western blot analysis using anti-ubiquitin antibody. A representative western blot for ubiquitinated JCN protein from one out of 3 different cell cultures is presented. (**D**) Neonatal mouse cardiomyocytes were incubated with palmitate or oleate in the presence of Murf1 inhibitor EMBL or vehicle. Upper panel: a representative western blot for JCN and bottom panel: quantification of JCN protein levels relative to GAPDH. (**E**) Neonatal mouse cardiomyocytes were infected with Ad-JPH2 or Ad-HA and then exposed to palmitate or oleate for 24 hours. Immunoprecipitation was performed to pull down Murf1 and its interacting proteins using anti-Murf1 antibody and western blot analysis determined the levels of JCN protein in Murf1 immunoprecipitates. Upper panel: representative western for JCN and Murf1 protein. Bottom panel: quantification of JCN/MURF1 protein in MURF1 immunoprecipitates. Data are mean ± SD, n = 3 different cell cultures. \**P*< 0.05 versus oleate + Ad-HA and #*P*< 0.05 versus palmitate + Ad-HA. (**F**) Neonatal mouse cardiomyocytes were infected with AD-shJPN2 or Ad-HA. Immunoprecipitation was performed to pull down Murf1 and its interacting proteins using anti-Murf1 antibody and western blot analysis determined the levels of JCN protein in Murf1 immunoprecipitates. A representative western from one out of 3 independent experiments for JCN and Murf1 protein shows knockdown of JPH2 increases the interaction between JCN and Murf1.

### Over-expression of JCN reduces cytosolic Ca^2+^ level and apoptosis in cardiomyocytes induced by stress or knockdown of JPH2

To determine whether JCN played a role in stress-induced cytosolic Ca^2+^ and apoptosis, we infected neonatal mouse cardiomyocytes with an adenoviral vector expressing JCN (Ad-JCN) or Ad-HA as a control and 24 hours later incubated them with palmitate, oleate, doxorubicin or saline for another 24 hours. Infection of Ad-JCN resulted in a higher level of JCN protein (**Figure S5**), lower level of cytosolic Ca^2+^ (**Figure 9A** and **9D)** and less caspase-3 activity and DNA fragmentation in palmitate- or doxorubicin-incubated cardiomyocytes as compared with infection of Ad-HA (**Figure 9B, 9C, 9E** and **9F**). Over-expression of JCN lowered JPH2 knockdown-elicited cytosolic Ca^2+^ level and inhibited apoptosis in neonatal mouse cardiomyocytes (**Figure 9G-9I**). In adult mouse cardiomyocytes, infection of Ad-JCN attenuated the higher cytosolic Ca^2+^ levels and cell death in response to palmitate (**Figure S6A-S6C**). The role of JCN was also examined in H9c2 cells. After transfection of a plasmid expressing JCN (pCMV-JCN) or EGFP (pCMV-EGFP) as a control, H9c2 cells were exposed to palmitate or oleate. Over-expression of JCN attenuated palmitate-induced higher cytosolic Ca^2+^levels and apoptosis in H9c2 cells (**Figure S7**).

**Figure 9.**
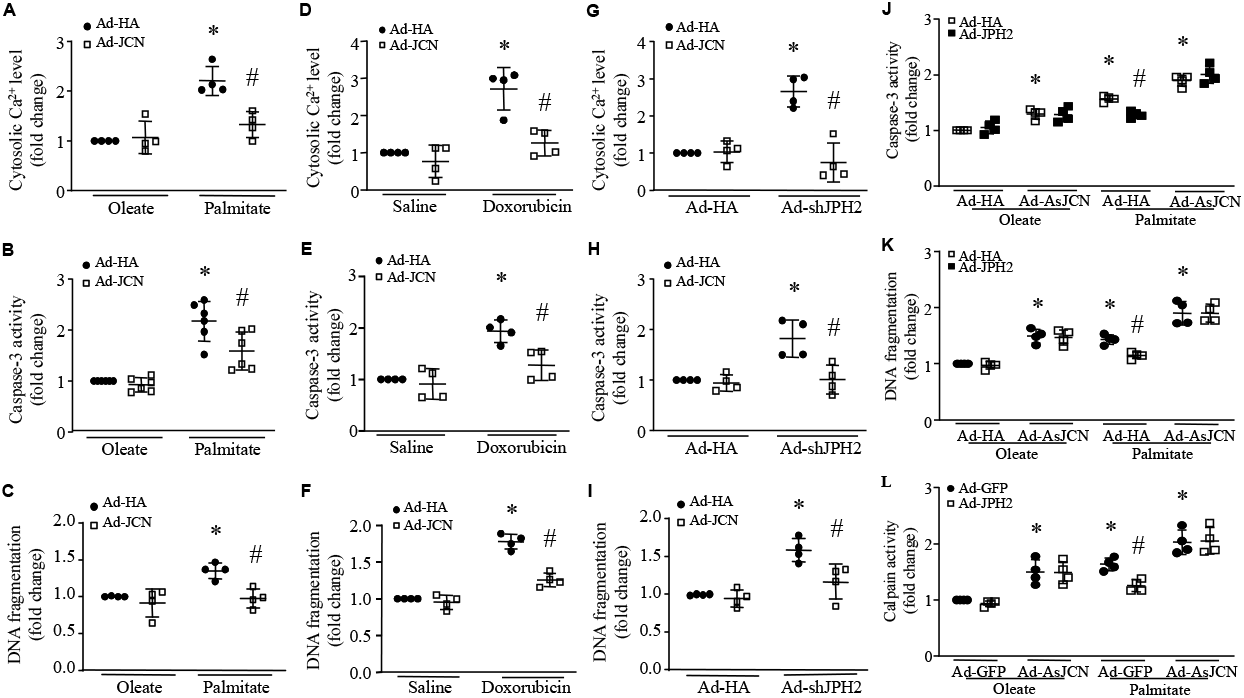
Role of JCN in cytosolic Ca^2+^ and apoptosis in cardiomyocytes under stress. (**A-C**) Neonatal mouse cardiomyocytes were infected with Ad-JCN or Ad-HA and then exposed to palmitate or oleate for 24 hours. Cytosolic Ca^2+^ (**A**), caspase-3 activity (**B**), and DNA fragmentation were analyzed (**C**). (**D-F**) Neonatal mouse cardiomyocytes were infected with Ad-JCN or Ad-HA and then exposed to doxorubicin or saline for 24 hours. Cytosolic Ca^2+^ (**D**), caspase-3 activity (**E**), and DNA fragmentation were analyzed (**F**). (**G-I**) Neonatal mouse cardiomyocytes were infected with Ad-JCN or Ad-HA in combination with Ad-shJPH2. Twenty-four hours later, cytosolic Ca^2+^ (**G**), caspase-3 activity (**H**), and DNA fragmentation were analyzed (**I**). (**J**-**L**) Neonatal mouse cardiomyocytes were infected with Ad-JPH2, Ad-HA or Ad-GFP in combination with an adenoviral vector containing *JCN* antisense (Ad-AsJCN), and then exposed to palmitate or oleate for 24 hours. Caspase-3 activity (**J**), DNA fragmentation (**K**), and calpain activity (**J**) were determined. Data are mean ± SD, n = 4-6 independent cell cultures. \**P*< 0.05 versus oleate + Ad-HA, saline + Ad-HA, Ad-HA + Ad-HA, oleate + Ad-GFP or oleate + Ad-HA, #*P*< 0.05 versus palmitate + Ad-HA, doxorubicin + Ad-HA, Ad-shJPH2+Ad-HA, palmitate + Ad-GFP or palmitate + Ad-HA.

### Knockdown of JCN offsets the effects of JPH2 over-expression on cytosolic Ca^2+^levels and apoptosis in cardiomyocytes under stress

To examine if JPH2 protected against palmitate-induced apoptosis through prevention of JCN protein degradation, we infected neonatal mouse cardiomyocytes with an adenoviral vector expressing anti-sense of JCN (Ad-AsJCN) or Ad-HA (or Ad-GFP) as a control, and then with Ad-JPH2 or Ad-HA (or Ad-GFP). Twenty-four hours after adenoviral infection, cardiomyocytes were incubated with palmitate or oleate for another 24 hours. Infection of Ad-AsJCN significantly lowered the JCN protein levels in cardiomyocytes (**Figure S8**). Similar to JPH2 knockdown, knockdown of JCN induced apoptosis determined as higher caspase-3 activity and DNA fragmentation in cardiomyocytes (**Figure 9J** and **9K**); however, JCN knockdown-elicited apoptosis was not prevented by JPH2 over-expression. Consistently, over-expression of JPH2 prevented palmitate-induced caspase-3 activity and DNA fragmentation in palmitate-induced cardiomyocytes; however, these effects of JPH2 over-expression were abolished in Ad-AsJCN infected cardiomyocytes where JCN was knocked down (**Figure 9J** and **9K**). These results indicate that the protective role of JPH2 is mediated through JCN in cardiomyocytes.

Since Ad-AsJCN concomitantly expressed GFP, which interfered with cytosolic Ca^2+^ measurement in cardiomyocytes, we analyzed calpain activity as an alternative indicator of cytosolic Ca^2+^ alteration as increased cytosolic Ca^2+^ induces calpain activation. Knockdown of JCN induced calpain activation, which was not inhibited by JPH2 over-expression (**Figure 9L**) whereas palmitate-increased calpain activity was attenuated by JPH2 over-expression in Ad-JPH2 infected cardiomyocytes. With JCN knockdown, over-expression of JPH2 was unable to attenuate the palmitate-stimulated calpain activity in cardiomyocytes. This result suggests that JCN is required for JPH2 maintaining cytosolic Ca^2+^ regulation in cardiomyocytes under stress.

## Discussion

The major findings of this study are that a reduction of JPH2 contributes to cardiomyocyte apoptosis under stress. Normal JPH2-JCN relationship protects against JCN ubiquitination and proteasome-dependent degradation through blocking Murf1-JCN interaction, thereby maintaining normal cytosolic Ca^2+^levels in cardiomyocytes. Further, over-expression of JCN attenuates apoptosis in cardiomyocytes under stress. This study reveals a novel role of JPH2 in stabilization of JCN and protection against cardiomyocyte death under stress.

JPH2 protein decreases in diseased hearts ^5–13^. Several mechanisms have been reported to modulate JPH2 expression at translational and post-translational levels including serum response factor-myocardin axis ^39^, miRNA-24 ^40^ and calpain activation ^41, 42^ as well as MMP2 ^43^. More recently, we found that aberrant autophagy is a new mechanism regulating JPH2 expression in cardiomyocytes ^35^. This study extended to report that both mRNA and protein levels of JPH2 are reduced in cultured cardiomyocytes under stress including exposure to palmitate and doxorubicin, indicating a regulation at mRNA levels. The reduction of *Jph2* mRNA expression is associated with aberrant autophagy as both palmitate and doxorubicin impairs autophagic flux and inhibition of autophagy with 3-MA prevents the reduction of *Jph2* mRNA levels in cardiomyocytes after exposure to palmitate and doxorubicin. However, it merits further investigation if the interplays between aberrant autophagy and other mechanisms are operative in regulation of JPH2 expression in cardiomyocytes under stress.

It is well known that JPH2 is required for the development, maturation and integrity of the t-tubule system, modulation of L-type Ca^2+^ channel and the gating stability of RyR2 in cardiomyocytes ^1, 4, 5, 14, 17–20^. A reduction of JPH2 may have consequences of Ca^2+^ uncoupling and abnormal Ca^2+^ leak, leading to disturbance of cellular Ca^2+^ homeostasis, which is known to cause cardiomyocyte dysfunction and death. In the current study, we extended the previous findings to demonstrate that JPH2 is required for cardiomyocyte survival as knockdown of JPH2 is sufficient to induced apoptosis in cardiomyocytes and that under stress a reduction of JPH2 is associated with apoptosis in cardiomyocytes. Notably, over-expression of JPH2 attenuates apoptosis in cultured cardiomyocytes under conditions of stress including palmitate, doxorubicin and hypoxia/re-oxygenation. Thus, the current study reveals an unappreciated role of JPH2 in preventing cardiomyocyte apoptosis under stress; and our data together with previous findings suggest that targeting JPH2 may be a useful therapeutic strategy to reduce cardiac pathology under stress. Indeed, transgenic over-expressionof JPH2 or its N-terminus,and gene therapy with its N-terminus attenuate myocardial remodeling and dysfunction in pre-clinical mouse models of heart failure ^12, 16, 44^.

An important finding of this study is that JPH2 is required for the stabilization of JCN in cardiomyocytes. Under stress, over-expression of JPH2 protected JCN against ubiquitination and proteasome-dependent degradation in cardiomyocytes. Several lines of evidence support this conclusion. First, conditions of stress decreased JCN protein level but not its mRNA levels. Second, knockdown of JPH2 resulted in lower protein levels of JCN in cardiomyocytes whereas over-expression of JPH2 preserved JCN protein in cardiomyocytes under stress. Third, incubation with proteasome inhibitor increased JCN protein levels in stressed cardiomyocytes. Fourth, conditions of stress increased JCN ubiquitination, which was prevented by JPH2 over-expression. Lastly, mass spectromtry revealed that lipid overload induced ubiquitination in mutiple residues of JCN protein and the non-ubiquitination mutant of JCN was resistant to lipid overload-induced degradation. Mechanistically, we found that stress increased the JCN-Murf1 interaction, which was prevented by JPH2 over-expression. Fruthermore, knockdown or pharmacological inhibition of Murf1 reduced JCN ubiquitination and increased JCN protein levels in cardiomyocytes under stress. Given that JPH2 had physical interaction with JCN, we propose a novel signaling mechanism contributing to the stabilization of JCN protein in cardiomoycytes: JPH2 binds to JCN and blocks the Murf1-JCN interaction thereby preventing Murf1-mediated ubiquitination and proteasome-dependent degradation of JCN. Currently, we can’t exclude such possibility that JPH2 brings de-ubiquitinating enzyme(s) to JCN. Nevertheless, specifically how JPH2 blocks Murf1-mediated ubiquitination of JCN requires further investigations.

Our finding that JPH2 protects JCN protein stability in cardiomoycytes has important relevance to cardiac patho-physiology as both have been implicated in maintaining Ca^2+^ homeostasis^5, 6, 22^. It is well known that JPH2 maintains proper Ca^2+^-induced Ca^2+^ release in cardiomyocytes^5, 6^. Notably, recent studies have demonstrated a role of JPH2 in stabilizing RyR2, which gates the release of Ca^2+^ from sarco/endoplasmic reticulum into the cytosol^14, 19, 20^., suggesting that JPH2 may function as a brake in Ca^2+^ release from RyR2, as loss of JPH2 results in aberrant Ca^2+^ leakage from the sarcoplasmic reticulum through RyR2^20^. Thus, it is most likely that a reduction of JPH2 may lead to more Ca^2+^ release from RyR2 into the cytosol, resulting in an elevation of cytosolic Ca^2+^, which contributes to apoptosis in cardiomyocytes under stress. In line with this, we showed that conditions of stress or knockdown of JPH2 resulted in higher levels of cytosolic Ca^2+^, which was attenuated by over-expression of JPH2 in cardiomyocytes. Furthermore, inhibition of RyR2 attenuated apoptosis in cardiomyocytes under stress. Although the mechanisms by which JPH2 stabilizes RyR2 in controlling Ca^2+^ release in cardiomyocytes remain largely unknown, our data suggest that prevention of JCN degradation may be one potential mechanism. Decreased JCN may increase RyR2 activity and consequent elevation of cytosolic Ca^2+^, which was reported to promote myocardial ischemia/reperfusion injury^24^. Notably, over-expression of JCN reduced stress-induced higher cytosolic Ca^2+^ in cardiomyocyte and knockdown of JCN abrogated the protections (inhibiting apoptosis and lowering cytosolic Ca^2+^) conferred by JPH2 over-expression in cardiomyocytes under stress, which argues that the role of JPH2 in stabilizing RyR2 may be mediated at least partially through JCN.

It is important to mention that using different types of cardiomyocytes (neonatal and adult mouse cardiomyocytes, and neonatal rat cardiomyocytes and H9c2 cells) and different conditions of stress (palmitate, doxorubicin and hypoxia/re-oxygenation), JPH2 exhibited similar protection against cardiomyocyte death and aberrant cytosolic Ca^2+,^ which strengths the notion that JPH2 is a survival factor of cardiomyocytes under both physiological and pathological conditions. However, there are some limitations in this study. First, in addition to Murf1, other enzymes in the process of JCN ubiquitination remain to be determined as ubiquitination involves three main steps: activation, conjugation, and ligation^45^. Second, we are aware of the difference of JCN (variant 1) protein sequence between human and mice (only 42% homology). Thus, future study will be needed to determine the regulation of JCN by JPH2 using human cardiomyocytes for translational/clinical significance.

In summary, we have provided evidence of a previously unrecognized role of JPH2 in maintaing JCN stability and subsequently preventing cardiomyocyte apoptosis under stress. This new role of JPH2 may be mediated by inhibiting Murf1-mediated JCN ubiquitination and proteasome-dependent degradation. Thus, both JPH2 and JCN may be new therapeutic targets for cardiomyocyte survival under physiological and pathological conditions.

## Author contributions

X.J., Y.H., R.N., and D.Z. conducted the experiments and analyzed the data; X.J., Y.H., G.C.F., D.L.J., L.S.S., Z.S. and T.P. designed the experiments and wrote the paper; G.C.F., D.L.J., L.S.S., S.C., Z.S. and T.P. discussed the data and revised the paper. All authors approved the final version to be published.

