## Supplementary Figures for "Junctophilin-2 promotes cardiomyocyte survival by blocking MURF1-mediated Junctin ubiquitination and proteasome-dependentdegradation"

### Supplementary Figure S1-S8

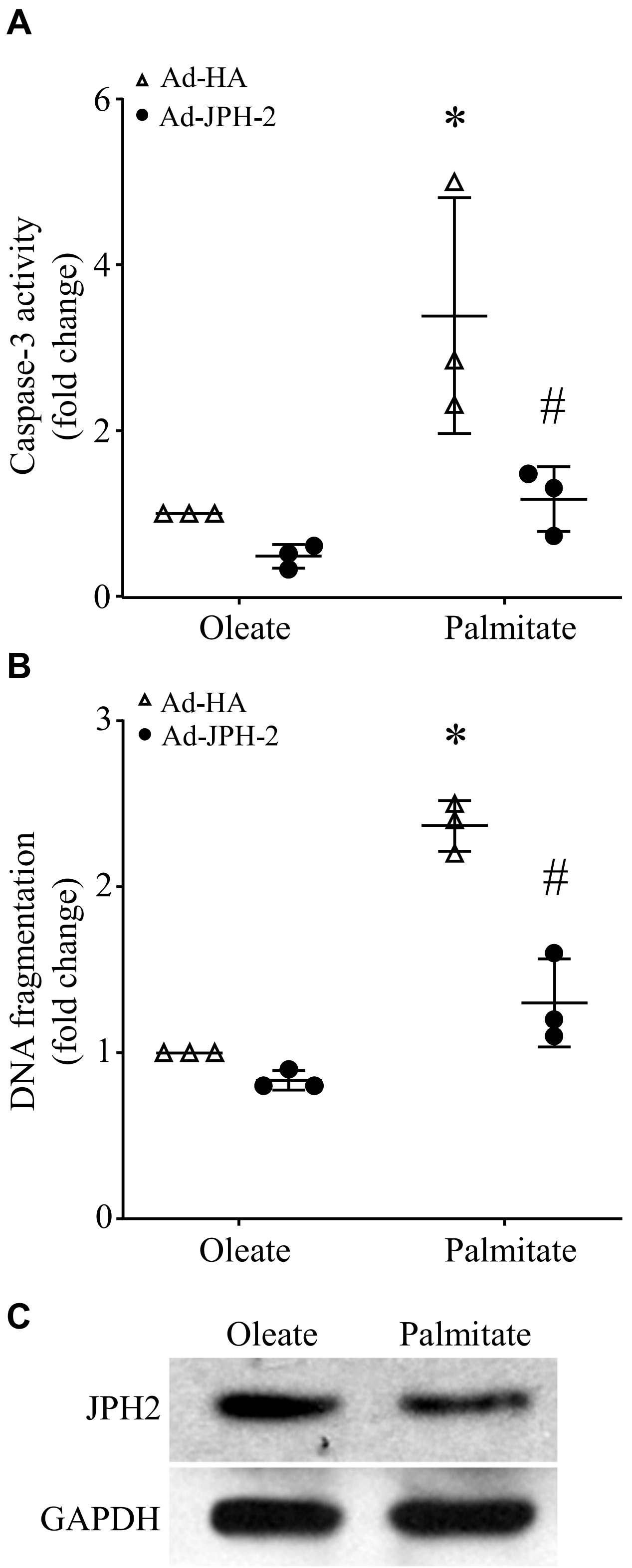


**Supplementary Figure S1. Effect of JPH2 over-expression on apoptosis in palmitate-induced neonatal cardiomyocytes**. Neonatal rat cardiomyocytes were infected with Ad-JPH2 or Ad-HA and then incubated with palmitate or oleate for 24 hours. Apoptosis was assessed by caspase-3 activity (A) and DNA fragmentation (B). (C) A representative western blot for JPH2 from one out of 3 independent experiments. Data are mean ± SD from 3 different culture cardiomyocytes. * $P < 0.05$ versus oleate + Ad-HA and # $P < 0.05$ versus palmitate + Ad-HA.

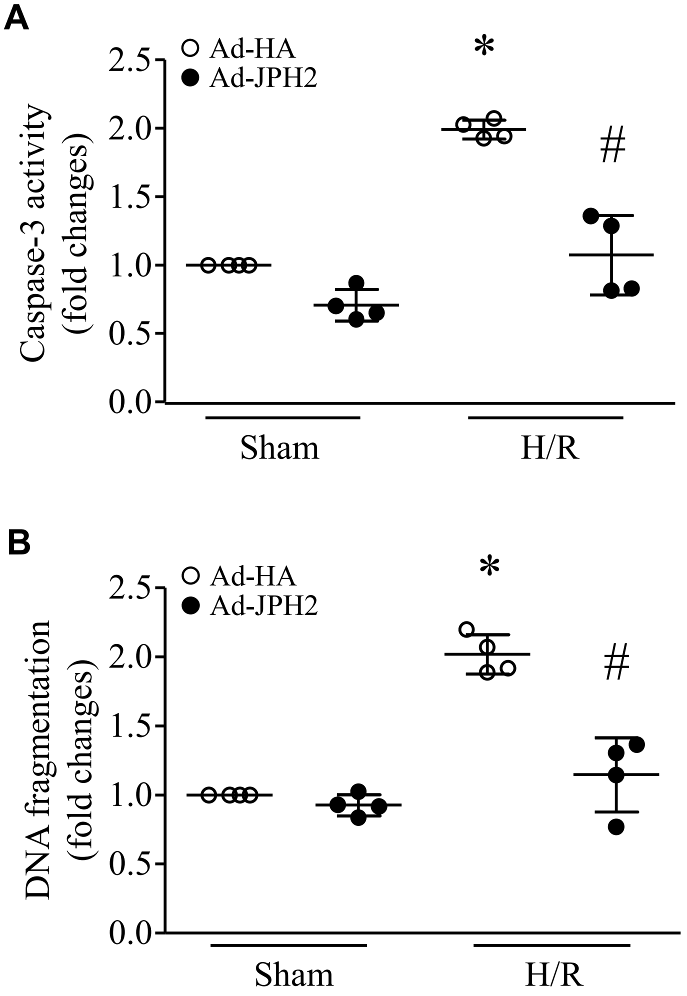


**Supplementary Figure S2. Role of JPH2 in apoptosis.** H9c2 cells were infected with Ad-JPH2 or Ad-HA and then subjected to a 24-hour period of hypoxia followed by re-oxygenation for another 24 hours (H/R) . Caspase-3 activity (A) and DNA fragmentation (B) were determined. Data are mean ± SD from 4 different cultures. * $P < 0.05$ versus sham + Ad-HA and # $P < 0.05$ versus H/R + Ad-HA.

| Peptide | Position | Score |
|---|---|---|
| ***MAEDKEAKHGGH | 5 | 2.59 |
| MAEDKEAKHGGHKNG | 8 | 1.25 |
| EAKHGGHKNGRRGGI | 13 | 2.26 |
| EGPGGLAKRKTKAKA | 86 | 3.10 |
| PGGLAKRKTKAKAKE | 88 | 3.99 |
| GLAKRKTKAKAKEPI | 90 | 3.50 |
| AKRKTKAKAKEPIKE | 92 | 2.57 |
| RKTKAKAKEPIKEEL | 94 | 3.55 |
| AKAKEPIKEELKKER | 98 | 2.95 |
| EPIKEELKKERGKAV | 102 | 2.24 |
| ELKKERGKAVPSKNE | 107 | 2.31 |
| RGKAVPSKNEERRQG | 112 | 1.93 |
| NEERRQGKKEQEDRG | 120 | 1.24 |
| EERRQGKKEQEDRGK | 121 | 1.06 |
| KEQEDRGKGRKKPDS | 128 | 0.82 |
| EDRGKGRKKPDSDTS | 131 | 3.03 |
| DRGKGRKKPDSDTSQ | 132 | 2.11 |
| PDSDTSQKASAAGKR | 140 | 1.61 |
| QKASAAGKRDRDKEK | 146 | 5.09 |
| AGKRDRDKEKASSDK | 151 | 1.63 |
| KRDRDKEKASSDKSS | 153 | 1.58 |
| KEKASSDKSSSKSKES | 158 | 3.76 |
| ASSDKSSSKSKESWKK | 161 | 2.17 |
| SDKSSSKSKESWKKAV | 163 | 2.73 |
| SKSKESWKKAVETKA | 167 | 3.24 |
| WKKAVETKAVSSKVA | 173 | 1.83 |
| ETKAVSSKVAARDKD | 178 | 2.98 |
| SKVAARDKDRRGRSS | 184 | 2.68 |
| SGHAHVSKENGQKRK | 199 | 1.72 |
| VSKENGQKRKN**** | 204 | 2.48 |
| KENGQKRKN****** | 206 | 4.18 |

**Supplementary Figure S3. Prediction of potential ubiquitination sites in JCN protein with Bayesian Discriminant Method**. Highlighted are ubiquitination modified residues in JCN protein from palmitate but not oleate-induced cardiomyocytes identified by mass spectrometry

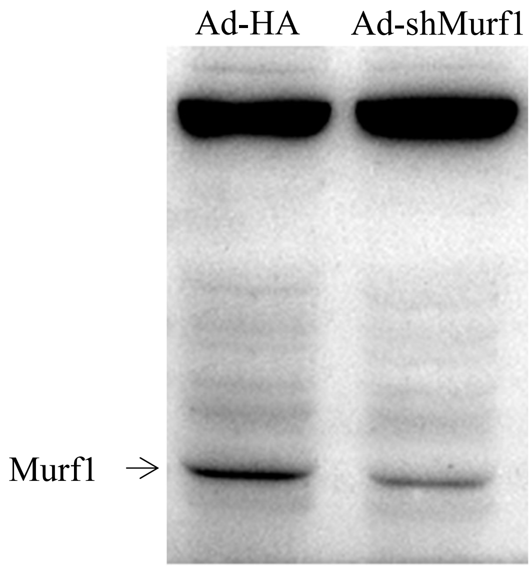


**Supplementary Figure S4. Knockdown of Murf1 in cardiomyocytes**. Neonatal rat cardiomyocytes were infected with Ad-shMurf1 or Ad-HA. Twenty-four hours later, the protein levels of Murf1 was determined by western blot analysis. A representative western blot shows that cardiomyocytes have lower levels of Murf1 protein after infection with Ad-shMurf1 compared with Ad-HA.

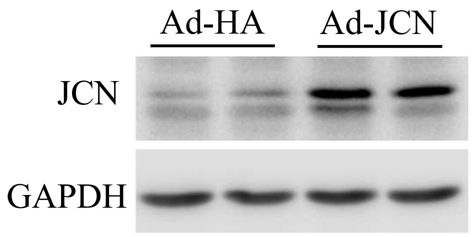


**Supplementary Figure S5**. **JCN protein levels in cardiomyocytes**. Neonatal mouse cardiomyocytes were infected with Ad-JCN or Ad-HA . Twenty-four hours later, the protein levels of JCN and GAPDH were analysed by western blot. A representative western blot for JCN and GAPDH is presented.

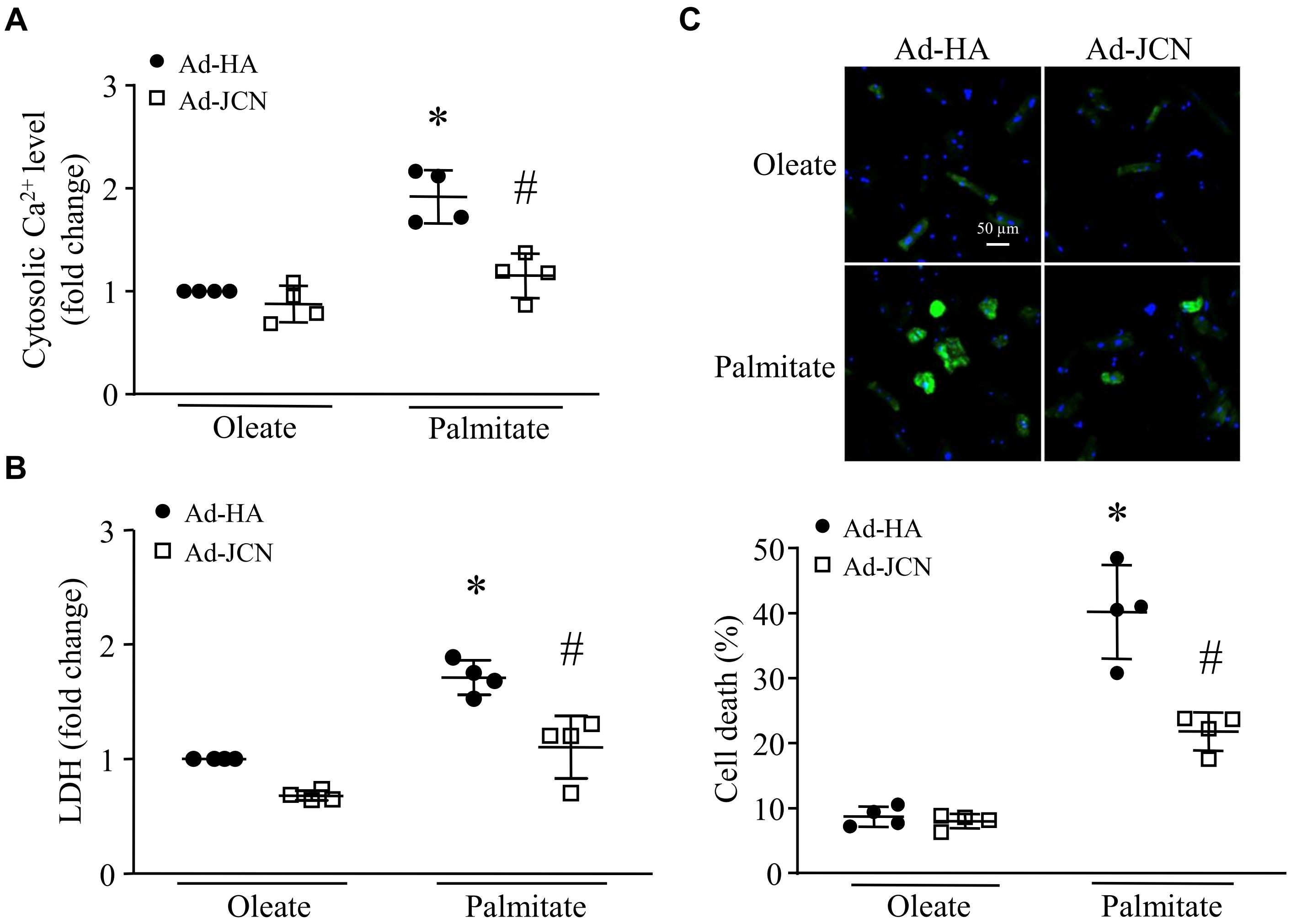


**Supplementary Figure S6**. Effects of JCN over-expression on cytosolic $Ca^{2+}$ and cell death. Adult mouse cardiomyocytes were infected with Ad-JCN or Ad-HA and then exposed to palmitate or oleate (0.2 mM) for 24 hours. (A) Cytosolic Ca2+ levels. (B) LDH release. (C) Upper panel: representative pictures for annexin V staining (green color for annexin V and blue color for nuclei) and bottom panel: quantification of annexin V staining positive cells. Data are mean ± SD, n = 4. *$P < 0.05$ versus oleate + Ad-HA and #$P < 0.05$ versus palmitate + Ad-HA.

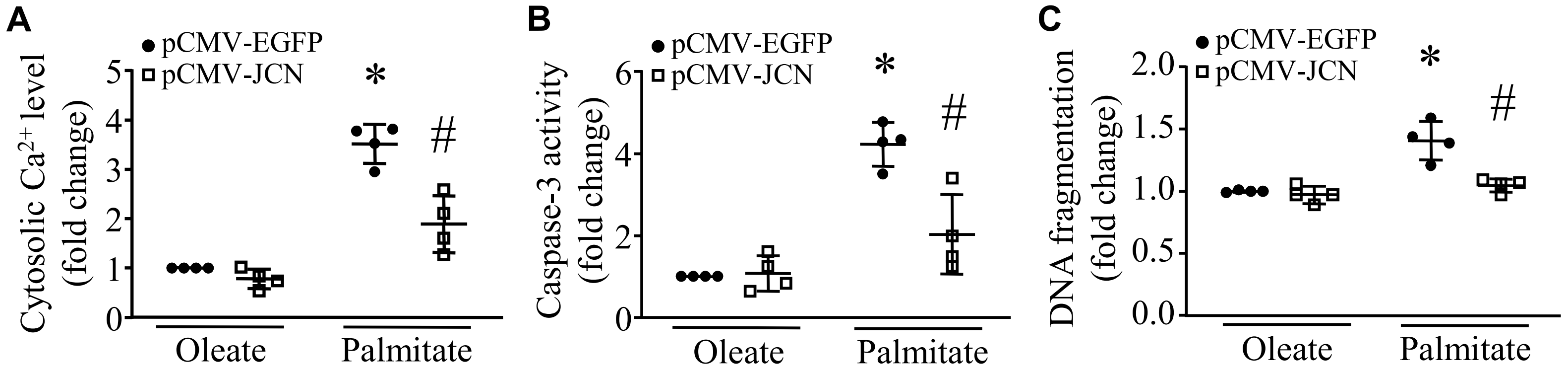


**Supplementary Figure S7**. **Effects of JCN over-expression on cytosolic $Ca^{2+}$ and apoptosis in cardiomyocytes**. H9c2 cells were transfected with plasmid DNA expressing JCN (pCMV-JCN) or expressing EGFP (pCMV-EGFP). Twenty-four hours later, H9c2 cells were exposed to palmitate or oleate (0.4 mM) for 24 hours. (A) Cytosolic $Ca^{2+}$ was determined. (B and C) Apoptosis was determined by caspase-3 activity and DNA fragmentation. Data are mean ± SD, n = 4 different cell cultures. * $P < 0.05$ versus oleate + pCMV-EGFP and # $P < 0.05$ versus palmitate + pCMV-EGFP.

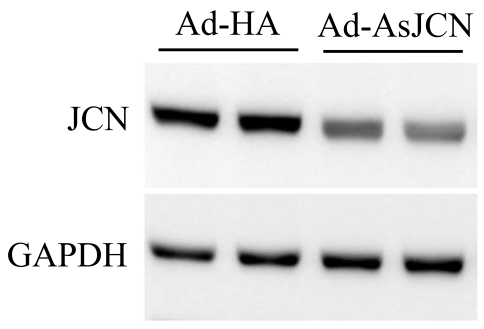


**Supplementary Figure S8**. **JCN protein levels in cardiomyocytes**. Neonatal mouse cardiomyocytes were infected with Ad-AsJCN or Ad-HA . Twenty-four hours later, the protein levels of JCN and GAPDH were analysed by western blot. A representative western blot for JCN and GAPDH is presented.
